# The GndA microprotein promotes cell growth during heat shock in *Escherichia coli*

**DOI:** 10.1101/2024.06.29.601336

**Authors:** Jessica J. Mohsen, Natalia R. Bertolotti, Kevin Jiang, Michael G. Mohsen, Christine M. DeRosa, Nikita Dewani, Laura Quinto, Georgina Dabdoub, Betel Erkalo, Ane Landajuela, Erdem Karatekin, Farren J. Isaacs, Sarah A. Slavoff

## Abstract

Hundreds of bacterial small open reading frames (sORFs) encoding microproteins of fewer than fifty amino acids have recently been identified. Biological functions have been ascribed to an increasing number of microproteins, some of which play roles in bacterial stress responses. In this work, we provide evidence that GndA, a frameshifted 36-amino acid microprotein encoded in an internal open reading frame overlapping the 6-phosphogluconate dehydrogenase (6PGD) coding sequence (CDS), functions in the response to heat shock. We demonstrate independent contributions of GndA and 6PGD to cell growth at high temperature. GndA is hydrophobic and associates with the pyruvate dehydrogenase complex and respiratory complex I (RCI). Consistent with a role in cellular homeostasis, GndA mutations alter the transcriptional response to heat shock. These results demonstrate that GndA promotes bacterial cell growth at high temperature independently of the 6PGD protein that it overlaps. GndA thus expands the paradigm of bacterial microproteins that function in cellular stress responses.

## Introduction

Due to application of a 50-amino acid cutoff for bacterial gene identification, bacterial sORFs encoding functional microproteins were overlooked until recently^1, 2^. This was because strategies for identification of longer, canonical proteins were not sufficiently sensitive to differentiate microprotein-coding vs. non-coding sORFs^3^. Microproteins are also difficult to detect using standard proteomic workflows^4^, and some microproteins exhibit limited conservation even among related species^5^. Despite these challenges, genomic^6, 7^, proteomic^4^ and molecular studies^18,19^ have now demonstrated that sORF-encoded microproteins (also referred to as miniproteins^8^, micropeptides^9^, or small proteins^10^) are numerous, functional and, in many cases, membrane-localized^11, 12^.

Bacterial microproteins have previously been reported to function in cellular stress responses^13^. For example, a foundational study reported upregulation of multiple microproteins in *Escherichia coli* during the heat shock response^11^. Additionally, predicted sORFs located in intergenic regions of *Enterobacteriaceae* were found to be expressed under Mg^2+^ starvation^14^, and the stressosome cytoplasmic complex activates the sigma B regulon when it interacts with the membrane-associated microprotein Prli42^8, 15^. Microproteins have also been implicated in mitochondrial ATP generation in eukaryotic cells. In human cells, mitochondrial microproteins regulate cellular respiration and assembly of the electron transport chain^16–19^. These studies position microproteins at the nexus of cellular metabolism and stress responses and suggest that microproteins may play regulatory roles in cellular homeostasis from bacteria to eukaryotes.

GndA is a 36-aa *E. coli* microprotein translated from a frameshifted, UUG-initiated sORF that internally overlaps the CDS of the metabolic enzyme 6-phosphogluconase dehydrogenase (6PGD), which functions in the pentose phosphate pathway (Figure 1A)^7, 20^. Both GndA and 6PGD are encoded within the *gnd* locus of *E. coli*. GndA was previously detected in the insoluble membrane fraction pelleted from *E. coli* cells subjected to heat shock and was predicted to form a transmembrane alpha helix^20^. However, the function of the *gndA* sORF has remained unexplored. We therefore sought to determine whether the GndA microprotein functions in the context of the bacterial heat shock response.

Internal, frameshifted sORFs (also termed intORFs) like GndA have been reported in bacteria and eukaryotes^21, 22^, but only a handful have been functionally characterized, and an evolutionary explanation for their existence is lacking. Providing a rationale for fitness benefit of internal sORFs, microproteins such as *comS* in *B. subtilis*^22–24^ and *rpmH* in *T. thermophilus*^25^ co-regulate phenotypes associated with the canonical CDS that they overlap. In this work, we report that GndA promotes cell growth at elevated temperature and transcriptional adaptation to heat shock, possibly as a result of its association with the pyruvate dehydrogenase complex or inner membrane proteins. These findings demonstrate that the overlapping ORFs encoded in the *gnd* gene locus, GndA and 6PGD, independently contribute to metabolic homeostasis and cellular fitness.

**Figure 1.**
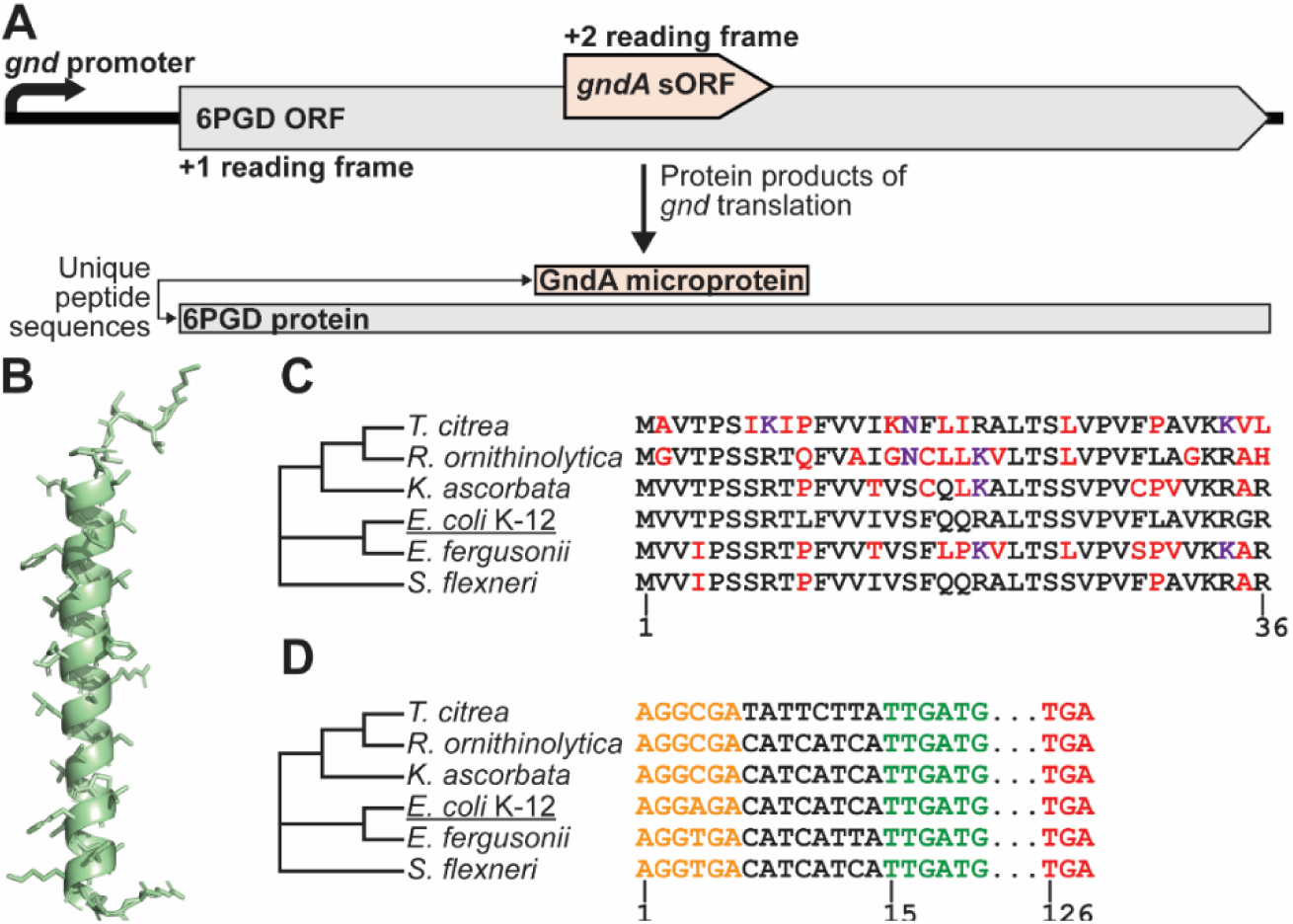
Evolutionary and structural analysis of the GndA microprotein. (*A*) The *E. coli* K-12 MG1655 *gnd* gene locus encodes two proteins in alternative reading frames with distinct amino acid sequences. Schematic of the *gnd* gene with the coding sequence (CDS) for 6PGD defined as the +1 reading frame. The GndA microprotein is encoded by an internal small open reading frame (sORF) in the +2 reading frame. As a result of alternative reading frame translation, GndA and 6PGD have completely different amino acid sequences. (*B*) GndA secondary structure prediction generated with AlphaFold 3. (*C*) GndA conservation in Gram-negative bacteria. Residues in black are identical, in purple are similar, and in red are not conserved. (D) The coding and translation regulatory sequences for GndA homologs were aligned. Shine Dalgarno sequence (orange), non-canonical TTG start codon (green), canonical ATG start codon (green), and stop codon (red) were manually identified.

## Materials and Experimental Details

### Evolutionary analysis

tblastn searches were run using the *E. coli* K-12 GndA amino acid sequence with standard parameters followed by manual inspection of start and stop codons. Nucleic acid sequences of top hits were aligned using Clustal Omega multiple sequence alignment.

### Bacterial genome editing

Scarless mutants at the *gnd* locus were generated with an SDS/colicin E1 selectable system^26^. WT *E. coli* K-12 MG1655 cells were transformed with a ColE1 kanamycin-resistant plasmid^27^ carrying anhydrotetracycline-inducible λ-Red recombineering genes (Exo, Beta, and Gam). To prepare cell cultures for gene editing, a single colony streaked out on LB agar with kanamycin was inoculated in LB broth with 50 µg/mL kanamycin at 37 °C and shaken at 200 rpm overnight.

The culture was diluted 1:100 in LB supplemented with 50 µg/mL kanamycin, and 100 ng/mL of anhydrotetracycline (Sigma catalog #37919) was added after 45 min of incubation to induce the λ-Red recombineering genes. Cultures were grown to OD_600_ = 0.5 and chilled on ice for 20 min. All tips, tubes, cuvettes, and autoclaved milli-Q water were chilled on ice. 1 mL of chilled culture was spun down at 3,000 rpm for 5 min at 4 °C and washed 3 times with chilled milli-Q water. To generate a Δ*tolC* strain from WT MG1655, an oligo mixture was prepared with 1 µL 100 µM oligonucleotide designed to disrupt the *tolC* locus added to 40 µL chilled milli-Q water. Washed cells were gently resuspended in oligo mixture and incubated on ice for 1 min. 50 µL of the cell-oligonucleotide mixture was transferred to a pre-chilled 1 mm electroporation cuvette (BioRad catalog #1652083) and transformed with 1.8 kV, 200 Ω, and 25 μF in a MicroPulser Electroporation Apparatus. Immediately after electroporation, cells were resuspended in 1 mL room temperature SOC medium (Sigma catalog #S1797-10X5ML). 2 mL LB Miller media was additionally added for a total recovery culture volume of 3 mL. Cells were recovered for a minimum of 2 h at 37 °C with 200 rpm shaking to allow turnover of TolC protein before selection. Recovery cultures were diluted 1:100 in LB supplemented with approximately 10 µg/mL colicin E1 (prepared as described previously^76^ in strain JC411) and 64 µg/mL vancomycin to select for *tolC* marker displacement, and incubated for 6-12 h at 37 °C with shaking at 200 rpm. Δ*tolC-* selected cultures were plated on antibiotic-free LB agar plates and individual colonies were screened for *tolC* knockout by both negative selection in LB supplemented with 0.005% SDS and colony PCR showing a lack of amplicon from the *tolC* locus.

All mutant strains, including Δ*tolC*, GndA^SPA^, ΔGndA^SCA^, ΔGndA^S116L^, ΔGndA^A117V^, 6PGD^A148V^, and Δ6PGD, were generated using sequential *tolC* selection/counterselection. At each target locus (*gnd*), the target gene was first knocked out in the MG1655 Δ*tolC* strain by knocking in the *tolC* cassette. This then served as a displaceable marker to select for knock in of tagged and/or mutant genes. First a Δ*gnd* strain lacking the *gnd* locus was constructed via double-stranded DNA (dsDNA) recombineering, replacing the native locus with *tolC*. (This strain was not used for experiments.) Recombineering cassettes were generated by amplifying the *tolC* cassette from MG1655 cells, with primers encoding 50-bp overhangs surrounding the gene targeted for deletion (Table S1). PCR products were purified by PCR purification (Qiagen catalog #28104) and full-length cassettes were subsequently isolated by agarose gel purification (Qiagen catalog #28506) and verified with Sanger sequencing. The Δ*tolC* strain bearing the recombineering plasmid was cultured and induced as described above, then electroporated with 1 µL of dsDNA substrate (100 – 400 ng of dsDNA). For positive *tolC* selection, cells were recovered for 2 h in SOC with LB, then plated on LB agar plates supplemented with 0.005% SDS. Individual knockout strains were confirmed for *tolC* integration by colony PCR and Sanger sequencing.

To generate tagged and mutant strains, the Δ*gnd* strain was subjected to another round of dsDNA recombineering to displace the *tolC* marker with the desired mutant sequence. Tagged and mutant cassettes (Table S1) were generated by colony PCR and overlap extension PCR with Phusion polymerase, or amplification from synthetic plasmids encoding the target sequence (Genscript), and confirmed by Sanger sequencing. The Δ*gnd* strain was cultured and induced as described above. Purified cassettes were electroporated into the strains, and cells were subjected to colicin E1 and vancomycin counterselection as in the initial *tolC* knockout experiment. After counterselection, mutant colonies were plated on LB agar and tested for desired mutations by negative selection in LB supplemented with 0.005% SDS as well as colony PCR and Sanger sequencing.

### Growth curve measurements

*E. coli* cell lines were streaked out on LB agar plates, or LB agar plates containing 25 µg/mL chloramphenicol when transformed with plasmids, and incubated 37 °C overnight. Single colonies were inoculated in LB broth and grown overnight at 37 °C with shaking at 200 rpm. Cultures were then diluted 1:100 and grown to exponential phase (OD_600_ = 0.3-0.6). Cultures were then normalized to OD_600_ = 0.2 and transferred to a sterile round bottom 96-well plate with lid. Three or four replicates per cell line were incubated at 45 °C with continuous orbital shaking while growth was monitored in a BioTek Synergy Neo2 Multimode Reader. OD_600_ readings were taken at 15 min intervals.

For transformation with plasmids (empty pBAD33 or pBAD33 encoding the GndA sequence without epitope tags) for rescue experiments, ΔGndA^SCA^, ΔGndA^S116L^, Δ*tolC*, and E. coli K-12 MG1655 cells were streaked out on LB agar plates and incubated at 37 °C overnight. Overnight cultures were inoculated from single colonies in LB broth with shaking at 200 rpm at 37 °C. Cultures were then diluted 1:100 in 7 mL of LB broth and grown to exponential phase (OD600 = 0.3-0.6). Cells were harvested by centrifugation at 3,000 rcf for 5 min at 25°C. Cells were then resuspended in 7 mL of LB broth and 1 mL aliquots were distributed into 1.5 mL microcentrifuge tubes. Cells were pelleted at 3,000 rcf for 5 mins at 4°C. Cells were then resuspended twice with 1 mL of ice-cold autoclaved Milli-Q water, pelleting at 3,000 rcf for 5 mins at 4°C following each wash. After the final wash, the pellet was resuspended in 1 mL ice-cold autoclaved 10% glycerol and again pelleted 3,000 rcf for 5 mins at 4°C. Plasmids were prepped in 50 μL of ice cold water to transform 1-10 ng of plasmid per cell line. The supernatant was discarded from the last wash and pellets were resuspended in their respective plasmid solutions and incubated on ice for at least 1 minute. Cells were then transferred to a MicroPulser Electroporation Cuvette with a 0.1 cm gap (BioRad, Catalog #1652083) and electroporated on the Ec1 setting (1.8 kV for 5 ms) using the BioRad MicroPulser Electroporator. Cells were immediately resuspended in 1 mL of room temperature S.O.C. media (Invitrogen, Catalog #15566-034), transferred to a new 1.5 mL microcentrifuge tube, and incubated for 1 h with shaking at 200 rpm at 37°C. Finally, cells were pelleted at 3,000 rcf for 5 min at 25°C, resuspended in 50 μL of LB broth and plated on LB agar plates supplemented with 25 µg/mL chloramphenicol. Growth curve measurements were carried out as above, except that a final concentration of 0.002% arabinose was added during growth to exponential phase.

### 6PGD quantification by targeted LC-MS/MS

Starter cultures were prepared by inoculating 5 mL of LB supplemented with 25 µg/ml chloramphenicol with a stab from a glycerol stock of each respective strain. Starter cultures were grown for approximately 16 hours at 37 °C with shaking at 200 rpm. 1 mL of starter culture was inoculated into 50 mL of LB supplemented as above. Cells were grown at 37 °C with shaking at 200 rpm until OD_600_ reached 0.6. Cells were harvested by centrifugation at 4000 rpm for 10 minutes at 4 °C and stored at -80 °C.

Cell pellets were resuspended in 1 mL of lysis buffer (50 mM Tris, 500 mM NaCl, 0.5 mM EDTA, 0.5 mM EGTA, 10% glycerol, and 1mL BugBuster Protein Extraction Reagent (Millipore 70921) and 1 cOmplete mini protease inhibitor tablet (Roche 11836153001) per 10 mL lysis buffer), incubated on ice for 30 min, and centrifuged at 15000 g for 10 min at 4°C to remove insoluble material. Chloroform/methanol precipitation was performed as previously described^28^. Precipitated protein pellets were redissolved in 200 µL of 8 M urea in 0.4 M Tris buffer (pH 8.0) and protein concentration was determined by UV absorbance spectroscopy at 280 nm on a nanodrop (Thermo Scientific ND-2000). 500 µg of protein was aliquoted, and volume was adjusted to 100 µL with 8M urea as above. 20 µL of 45 mM DTT was added to each sample and the reaction was incubated at 60°C for 30 min in a water bath and then quenched on ice for 30s. 4 µL of 100 mM iodoacetamide (IAA) solution was added and alkylation of cysteines proceeded for 30 minutes at room temperature in the dark. Excess IAA was quenched with 4 µL of 45 mM DTT. Samples were diluted with 1 mL water followed by the addition of 33.3 µg of sequencing grade modified trypsin (Promega V5111). Trypsin digestion was performed at 37°C overnight. After incubation, 10 µL of LC-MS grade TFA (Thermo Scientific 85183) was added to acidify samples and pH was tested to be below 3 using pH test paper (Fisherbrand™ 13-640-506). Acidified samples were desalted using Sep-Pak C18 Vac Cartridges (Waters WAT023590).

Samples were resuspended in 50 µL 0.1% FA, followed by centrifugation at 15,000 rcf at 4°C for 30 min. A 2 µL aliquot of each sample was injected onto a pre-packed column attached to a Vanquish Neo UHPLC (Thermo Scientific) in-line with a Thermo Scientific Orbitrap Eclipse Tribrid mass spectrometer. Samples were analyzed with a 120 min gradient as follows (solvent A: 0.1% FA; solvent B: acetonitrile (ACN) with 0.1% FA): (min/%B) 0/6.0, 90/27.0, 100/45.0, 110/100.0, 120/100.0. The full MS was collected using a data-dependent acquisition method over the mass range of 350-1400 m/s with a resolution of 120,000 and AGC target of 4 × 10^5^. Ions were isolated in quadropole mode with a 1.4 m/z isolation window and fragmented with higher-energy collisional dissociation (HCD) at 30% collision energy. Ions were detected at the orbitrap detector with 30,000 resolution and dynamic exclusion was 20s. Files were analyzed using MaxQuant, and oxidation of methionine and N-terminal acetylation were set as variable modifications. *E. coli* K-12 Uniprot plus GndA was used as the database for searching.

### Protein extraction and co-immunoprecipitation (co-IP) and LC-MS/MS

For protein extraction from GndA^SPA^ cells, single colonies were inoculated in LB broth and grown overnight with shaking at 200 rpm at 37 °C. Cultures were diluted 1:100 in 10 mL of fresh LB broth and grown to exponential phase (OD_600_ = 0.3-0.5). Exponential cultures were then grown for 2 hours at 37 °C or 45 °C, and were then immediately collected for protein extraction as described.

Equal numbers of cells normalized by OD_600_ were taken from normal (37 °C, 2 hours) and heat shocked cultures (45 °C, 2 hours) and pelleted by centrifugation at 21 °C at 4,000 rpm for 5 min. Media was decanted and pellets were resuspended in 1 mL of B-PER lysis buffer (Thermo Scientific catalog# 90078) containing DNase I, Lysozyme, and Roche complete protease inhibitor (Sigma catalog #11836170001). Cells were lysed by sonication with a QSonica Misonix Microson Ultrasonic Cell Disruptor XL-2000 for 5 rounds, 5 s on 25 s off, on ice at medium (5) setting. Lysate was clarified by centrifugation at 4 °C at 21,000 rcf for 5 min. Clarified lysate containing solubilized protein was removed without disrupting pelleted material and transferred to a fresh tube.

For each clarified lysate, 25 µL of anti-FLAG M2 affinity agarose gel (Sigma catalog #A2220) was washed in 1 mL ice-cold B-PER and collected at 1,000 rcf for 1.5 min at 4 °C. Wash supernatant was removed without disrupting agarose gel, and the protein containing clarified lysate was added. The protein lysate was incubated with agarose gel at 4 °C with rotation for 1 h or overnight. Samples were spun down at 1,000 rcf for 1.5 min at 4 °C to collect beads bound to FLAG fusion protein and binding partners. To remove nonspecific binding, supernatant was removed and beads were washed twice with TBS containing 1% Triton X-100 (Sigma catalog #T8787) and 500 mM NaCl at 4 °C with rotation for 3 min. A final wash with 10 inversions by hand was done in TBS containing 1% Triton X-100 and 150 mM NaCl to return proteins to physiological salt concentration. Bound protein was eluted off agarose gel with 30-50 μL 1X 3XFLAG peptide (Sigma catalog #F4799) at 4 °C with rotation for 1 h or boiled. Agarose beads were collected at 4 °C 1,000 rcf for 2 min, and supernatant containing immunoprecipitated protein was removed and stored at -80 °C for Western blotting to confirm successful co-IP of GndA^SPA^ and for subsequent LC-MS/MS.

Quantitative proteomics was performed as previously reported^29^. Gel slices containing entire lanes were digested with trypsin at 37 °C for 14-16 h. The resulting peptide mixtures were extracted from the gel, dried, subjected to ethyl acetate extraction to remove residual detergent, de-salted with a peptide cleanup C18 spin column (Thermo Scientific catalog #89870), then resuspended in 35 µL 0.1% formic acid (FA), followed by centrifugation at 15,000 rcf at 4 °C for 30 min. A 5-µL aliquot of each sample was injected onto a pre-packed column attached to an Easy-nLC 1200 (Thermo Scientific) in-line with a Thermo Scientific Q Exactive Plus Hybrid Quadrupole-Orbitrap mass spectrometer. A 125-min gradient was used to further separate the peptide mixtures as follows (solvent A: 0.1% FA; solvent B: 80% acetonitrile (ACN) with 0.1% FA): (min/%B) 0/5%, 90/45%, 91/85%, 101/85%, 102/5%, 125/5%. The full MS was collected over the mass range of 300-1,700 m/z with a resolution of 70,000 and the automatic gain control (AGC) target was set as 3 × 10^6^. MS/MS data was collected using a top 20 high-collisional energy dissociation method in data-dependent mode with a normalized collision energy of 28.0 eV and a 1.6 m/z isolation window. MS/MS resolution was 17,500 and dynamic exclusion was 90 s. Files were analyzed using MaxQuant, and oxidation of methionine and N-terminal acetylation were set as variable modifications. *E. coli* K-12 Uniprot plus GndA was used as the database for searching. Protein quantitation was accomplished with MaxQuant LFQ (version 2.0.2.0)^30^.

### Protein extraction and co-immunoprecipitation (co-IP) followed by Western blotting

GndA^SPA^ cells transformed with pBAD33 plasmids encoding potential interaction partners were grown on LB agar with 25 µg/mL chloramphenicol. Overnight cultures were inoculated from single colonies in LB broth with shaking at 200 rpm at 37 °C. Cultures were diluted 1:100 in 20 mL of fresh LB broth and grown to exponential phase (OD_600_ = 0.3-0.5). Exponential cultures were then induced with 40 µL of 10% arabinose per sample for a final concentration of 0.02% arabinose, and then grown for 2 hours at 45 °C. Samples were immediately collected for protein extraction.

Equal numbers of cells, normalized by optical density (OD_600_), were harvested by centrifugation at 4,000 rpm for 5 min at 4°C. Cell pellets were resuspended in 1 mL B-PER bacterial protein extraction reagent (Thermo Scientific, catalog #90078) supplemented with DNase I, lysozyme, and protease inhibitor cocktail (Roche Complete; Sigma-Aldrich, catalog #11836170001). Cells were lysed by sonication (Microson, Model no: XL2000 ultrasonic cell disrupter, 10 seconds on, and 30 seconds off, for 3 cycles at 7 Watts), and lysates were clarified by centrifugation at 15,000 rcf for 10 min at 4 °C. The clarified supernatant containing soluble proteins was transferred to fresh microcentrifuge tubes.

For co-immunoprecipitation of SPA-tagged GndA, clarified lysates were incubated with Pierce Anti-DYKDDDDK (FLAG) Magnetic Agarose Beads (Cat # A36797, Thermo Scientific™). 50 µL of anti-FLAG magnetic beads were used per 480 µL of clarified lysate. Prior to use, beads were equilibrated by washing three times with B-PER buffer using a magnetic stand and then half the beads were pre-incubated with 50 uL of 50X 3XFLAG peptide (Cat #F4799, Sigma) in 400 µL of B-PER for 2 hours at 4 °C. Lysates were then incubated with either pre-blocked or equilibrated beads at 4 °C with gentle rotation overnight. Following incubation, beads were collected using a magnetic separator and subjected to sequential washes to reduce non-specific binding. An initial high-stringency wash using buffer supplemented to 500 mM NaCl, followed by a final wash at 150 mM NaCl. Beads were washed a total of three to five times. Bound proteins were eluted by boiling beads in SDS sample buffer with β-mercaptoethanol at 95–100 °C for 5–10 min and analyzed by SDS-PAGE or Tricine gel followed by Western blotting as described below.

### Western blotting

Standard Western blotting procedures were followed with modification using a Mini-PROTEAN Tetra Cell, Mini-Trans blot module, and power supply (Bio-Rad catalog# 1658033). Protein samples were resolved on 16% Tricine-SDS-PAGE for proteins with molecular weights less than 20 kDa^31^, and 10% or 15% Glycine-SDS-PAGE was used for proteins with molecular weights above 20 kDa. 4X SDS loading buffer was added to protein samples extracted from equivalent *E. coli* samples. PAGE was run at 50-80 V as proteins migrated through the stacking gel, and at 70-90 V through the resolving gel until the Protein Xtra Prestained Ladder (Bio-Rad catalog # 1610377) achieved good separation. Gels were equilibrated in pre-chilled transfer buffer with 15% methanol and then protein was transferred to a 0.2 µm nitrocellulose membrane (Bio-Rad catalog# 1620112) for proteins with molecular weights less than 20 kDa, or a 0.45 µm nitrocellulose membrane (Sigma catalog# 10600002) for proteins with molecular weights above 20 kDa, via wet electroblotting on ice for 1 h at 300 mV. Membrane was blocked in 3% BSA in TBS-T for 2 h with rocking at room temperature. Primary antibody was added 1:1,000 in 3% BSA in TBS-T and incubated at room temperature for 1 h with rocking, or overnight at 4 °C with nutation. The membrane was washed three times in TBS-T for 5 min at room temperature with rocking. Secondary antibody was added 1:10,000 in 3% BSA in TBS-T and incubated 1 h at room temperature with rocking. Three washes in TBS-T were performed. Chemiluminescent imaging was done with horseradish peroxidase conjugated IgG and visualized with Clarity western ECL substrate (BioRad catalog# 1705061). Images were taken on BioRad ChemiDoc XRS+ Gel Imaging System.

### Antibodies for Western blotting

DYKDDDDK tag rabbit monoclonal antibody (Cell Signaling catalog #14793S), anti-FLAG mouse M2 monoclonal antibody (Sigma Aldrich catalog #F1804), Myc tag rabbit polyclonal antibody (Rockland catalog #600-401-381), HRP-conjugated secondary antibodies (anti-mouse or anti-rabbit) (Rockland catalog #611-1302)

### RNA-seq

Δ*tolC*, ΔGndA^SCA^ and ΔGndA^S116L^ strains were streaked on LB agar plates and incubated 37 °C overnight. Single colonies were inoculated in LB broth and grown overnight with shaking at 200 rpm at 37 °C. Exponential cultures were then grown for 2 hours at 45 °C. RNA extraction was carried out using the RNeasy Protect Bacteria kit (Qiagen catalog #74524). 5 × 10^8^ *E. coli* cells were harvested from each sample using an RNeasy mini spin column. Qiagen protocol 4 was followed for enzymatic lysis with lysozyme (DOT Scientific catalog #DSL38100) and 20 µL proteinase K (NEB catalog #P8107S). Qiagen protocol 7 was followed for RNA purification including DNase I digestion on-column (Qiagen catalog# 79254). Total RNA was eluted in 50 µL RNase free water.

Each strain was analyzed in biological triplicate. For each sample, 2 μg of extracted RNA per replicate was submitted to the Yale Center for Genome Analysis, where ribosomal RNA depletion and library preparation was performed. Sequencing was performed with the Illumina NovaSeq system at a depth of approximately 20 million reads per sample. Paired-end reads were sequenced with a read length of 150 base pairs.

Raw sequencing reads were assessed for quality using FastQC (v0.12.1) (https://www.bioinformatics.babraham.ac.uk/projects/fastqc/) and trimmed to remove adapter sequences and low-quality bases using cutadapt (v5.2). Ribosomal RNA contamination was reduced by filtering reads against *Escherichia coli* rRNA sequences using BBDuk (v39.52). Filtered reads were aligned to the *E. coli* K-12 MG1655 reference genome (NCBI assembly GCF_000005845.2) using Bowtie2 (v2.5.2). Alignment files were converted to sorted and indexed BAM format using SAMtools (v1.22.1), from which gene-level read counts were generated using featureCounts (v2.1.1) with the *E. coli* K-12 MG1655 annotation and summarization at the gene level. Quality control metrics across all processing steps were aggregated using MultiQC (v1.28).

Downstream analysis was performed in R (v4.4.1). Raw count matrices were filtered to remove lowly expressed genes, retaining genes expressed in at least 50% of samples and with a minimum total count threshold of 10. Differential expression analysis was conducted using the DESeq2 package (v1.44.0), which models count data using a negative binomial generalized linear model and incorporates normalization for sequencing depth and dispersion across biological replicates. The analysis used a model that included batch as a covariate alongside genotype to account for technical variation between sequencing runs. Differential expression was assessed by comparing each mutant to the Δ*tolC* background. *P*-values were derived from Wald tests and corrected for multiple testing using the Benjamini–Hochberg method. Genes with an adjusted *p*-value < 0.05 and an absolute log2 fold change > 1 were considered significantly differentially expressed and were used in all subsequent analyses. Genes significantly upregulated in both mutants were subjected to Gene Ontology (GO) analysis after removing y-ome genes^32^.

## Results and Discussion

### Results

In prior work, we provided proteomic and genetic evidence for expression of GndA from an internal, frameshifted sORF overlapping the 6PGD coding sequence^20^. We showed that GndA appears in the insoluble membrane fraction of *E. coli* cells during heat shock. In a subsequent report by Mankin and colleagues, ribosome profiling confirmed translation of GndA from a non-canonical UUG start codon^7^. We hypothesized that GndA plays a role in the response to elevated temperature in *E. coli* and utilized the Mankin group’s identification of the full-length GndA sequence to reanalyze its properties and interrogate its function. We first re-examined its predicted structure. Many microproteins exhibit single transmembrane helices^33–35^, and AlphaFold 3^36^ predicts the GndA peptide to form an alpha helix extending from residues 7 to 35 (Figure 1B).

While not all adaptive microproteins are conserved^37^, conservation remains a robust metric to identify microproteins with phenotypic effects^38^. We thus hypothesized that if homologs are present in multiple bacterial lineages, GndA is more likely to play a role in cellular fitness. To examine GndA conservation, we first performed a translated nucleotide BLAST (tblastn) search using the *E. coli* GndA microprotein sequence. We then retrieved the genomic DNA sequences of putative homologs and manually identified the Shine Dalgarno sequence, start and stop codon in each. 23 GndA homologs with > 50% sequence similarity were identified in Gram-negative bacteria using these criteria, all found in syntenic sORFs internally overlapping the 6PGD CDS of the respective organism (Figure 1C-D, Figure S1). While the conclusions from this analysis are limited by evolutionary constraint imposed by selection on the overlapping 6PGD reading frame, the presence beyond *E. coli* of homologous GndA sequences could be consistent with functionality.

We created a series of mutant *E. coli* K-12 MG1655 strains to examine the function of GndA in heat shock (Figure 2A-E, top panels, and Tables S1 and S2). Three different strategies to abrogate GndA expression were pursued. To eliminate GndA translation without perturbing 6PGD sequence or expression, a start codon ablation mutant (ΔGndA^SCA^) was generated in which the validated start site (UUG) and two additional in-frame AUG codons were mutated (Figure 2A). All mutations in the ΔGndA^SCA^ strain are silent in the 6PGD reading frame. In addition, we created mutant strains in which nonsense mutations prematurely truncating the GndA sORF were introduced. Because we were unable to identify positions at which nonsense mutations could be introduced in the GndA reading frame without altering the 6PGD amino acid sequence, we selected two mutations we judged to be most conservative with respect to 6PGD while truncating GndA early enough to likely result in loss of function. By this logic, strains ΔGndA^S116L^ and ΔGndA^A117V^, in which premature stop codons were introduced after residues 17 and 18 of GndA, were generated (Figure 2B-C). ΔGndA^S116L^ and ΔGndA^A117V^ introduce S116L or A117V point mutations in the 6PGD coenzyme domain, respectively, which are reflected in the strain names. For comparison, strains with mutations in 6PGD only were created, first by inducing a point mutation in the same domain of 6PGD but downstream of the *gndA* sORF (6PGD^A148V^) (Figure 2D), and second by deleting the 6PGD start site (Δ6PGD) (Figure 2E). The GndA sORF is not perturbed in either of these 6PGD mutant strains. We note that all mutants used in this study lack *tolC*, which was deleted as a selection marker for our recombineering strategy^26^. We therefore utilized an *E. coli* K-12 MG1655 *tolC* deletion strain (Δ*tolC*) as the isogenic control for phenotypic experiments, though we note that Δ*tolC* cells grew comparably to wild-type *E. coli* K-12 at both 37 and 45 °C (Figure S2).

**Figure 2.**
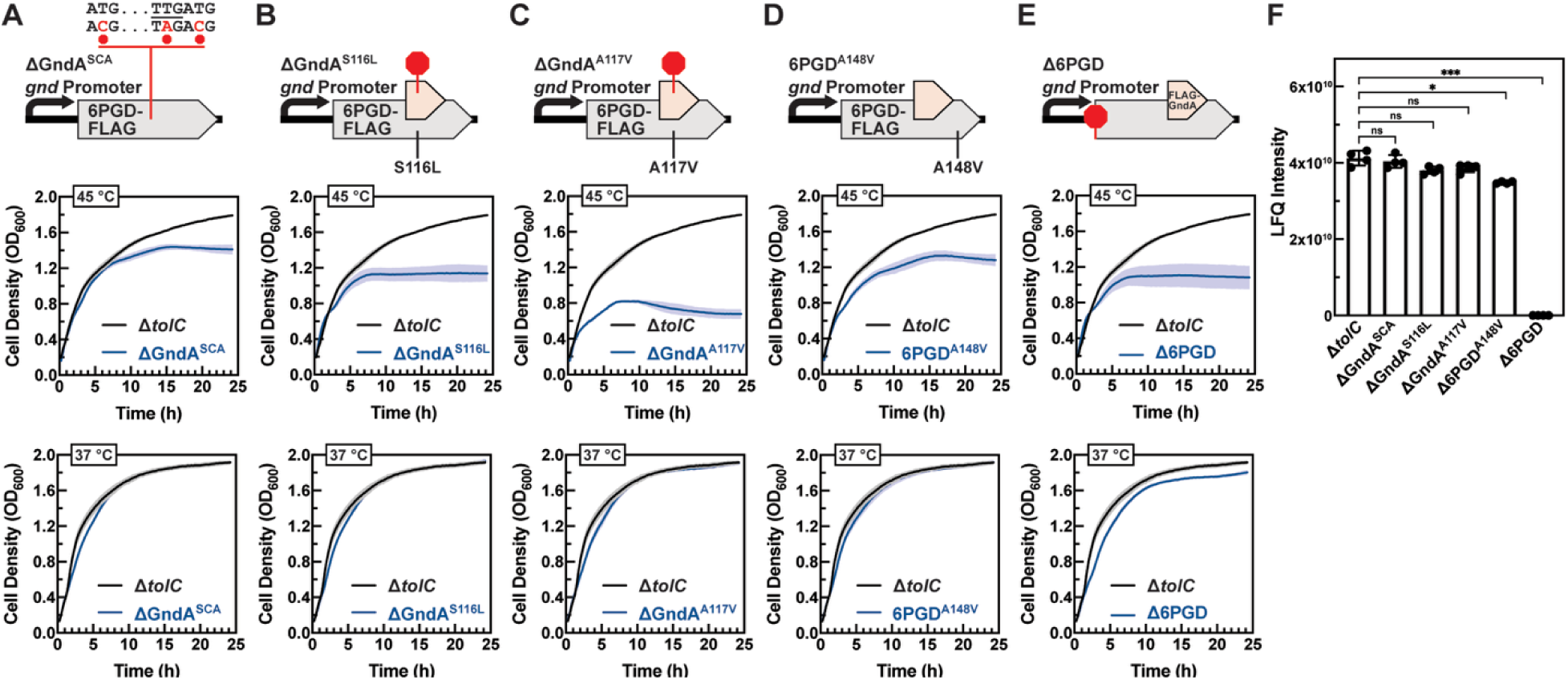
GndA mutants exhibit a reduced growth phenotype at elevated temperature. (A-E) Top, schematic of genomic edits in each cell line bearing (A) mutations in the GndA start codons that are silent in the 6PGD reading frame, (B-C) a truncating mutation in GndA creating a point mutation in 6PGD, or (D-E) mutation or deletion of 6PGD alone. Middle, growth curves (OD_600_) for mutant cell lines (blue) and controls (black) bearing the same Δ*tolC* genetic background, grown at 45 °C. Bottom, growth curves (OD_600_) for mutant cell lines (blue) and controls (black) bearing the same Δ*tolC* genetic background, grown at 37 °C. Error (shading above and below curves) calculated from four replicates. (F) Label-free proteomic quantitation (LFQ) of 6PGD protein levels was carried out in Δ*tolC* control and experimental mutant strains used in (A-E). Bars represent the mean of four replicates. Error, s.d.; significance, Student’s *t*-test. ns, not significant; *, *p*-value < 0.05; ***, *p*-value < 0.001.

We then examined whether GndA plays a role in cell growth at elevated temperature. Little difference was observed in growth rates of the control Δ*tolC* cells and the mutant strains when grown at 37 °C (Figure 2A-E, bottom panels). ΔGndA^SCA^ cells exhibit a reduced cell density during stationary phase at 45 °C relative to Δ*tolC* cells (Figure 2A, middle panel). Supporting this finding, GndA mutants bearing premature stop codons (ΔGndA^S116L^ and ΔGndA^A117V^) exhibited comparable, or more pronounced, growth defects at high temperature (Figure 2B-C, middle panels). As expected, 6PGD expression and activity also contribute to fitness at elevated temperature: introduction of a point mutation in the coenzyme domain downstream of the *<u>gndA</u>* sORF (6PGD^A148V^) or deletion of its start codon (Δ6PGD) caused reduced density at stationary phase (Figure 2D-E, middle panels). In order to unambiguously attribute the growth defects observed in the GndA mutant strains to loss of GndA, and exclude possible effects on expression or activity of 6PGD, we quantified 6PGD protein levels across all mutant and control strains using shotgun proteomics with label-free quantitation (Figure 2F). 6PGD levels in the ΔGndA^SCA^, ΔGndA^S116L^ and ΔGndA^A117V^ strains were unchanged compared to Δ*tolC* cells, while a small, statistically significant decrease in 6PGD abundance was observed in 6PGD^A148V^. 6PGD expression was undetectable in Δ6PGD cells, as expected. We emphasize that the ΔGndA^SCA^ cell line completely abrogates translation of GndA but introduces no mutations in the 6PGD reading frame, and exhibits 6PGD expression levels equal to controls. We can therefore conclusively attribute the growth defect observed in ΔGndA^SCA^ cells to the absence of GndA. The observed growth defects in ΔGndA^S116L^ and ΔGndA^A117V^ strains may be due in part to the premature truncating mutations in GndA, but we cannot exclude the possibility that the point mutations in 6PGD also contribute to the phenotype observed. GndA and 6PGD thus independently promote *E. coli* cell growth at 45 °C.

To further support the specific contribution of GndA to cell growth at elevated temperature, we “rescued” the ΔGndA^SCA^ and ΔGndA^S116L^ strains with GndA encoded on a pBAD33 arabinose-inducible vector, or the empty pBAD33 vector as a control. We observed that the mutant strains carrying any pBAD33 vector exhibit impaired growth compared to wild-type *E. coli* K-12 MG1655 or Δ*tolC* cells at both 37 and 45 °C, even in the absence of arabinose (Figure S3), which we hypothesize could be due to greater sensitivity of these strains to cellular stress associated with plasmid DNA electroporation. Due to these growth defects, which appear to be largely irreversible, we found that mutant cell lines carrying rescue plasmids could not be stored as glycerol stocks. We therefore freshly transformed the plasmids for each rescue experiment. We observed a very small but reproducible improvement in 45 °C growth relative to empty pBAD33 vector control for the ΔGndA^SCA^ strain, though this did not always rise above the variation of our measurements; we observed a more substantial increase in growth in the ΔGndA^S116L^ strain (Figure 3). There was no effect of rescue at 37 °C for either strain (Figure S4). The blunted rescue at high temperature may be a consequence of the severe growth defect induced by plasmid introduction, and the mechanistic basis of the observed strain-specific differences in rescue efficacy remains to be determined. Even given these limitations, the partial rescue observed, particularly in the ΔGndA^S116L^ strain, is consistent with a beneficial role for GndA in *E. coli* growth at 45 °C.

**Figure 3.**
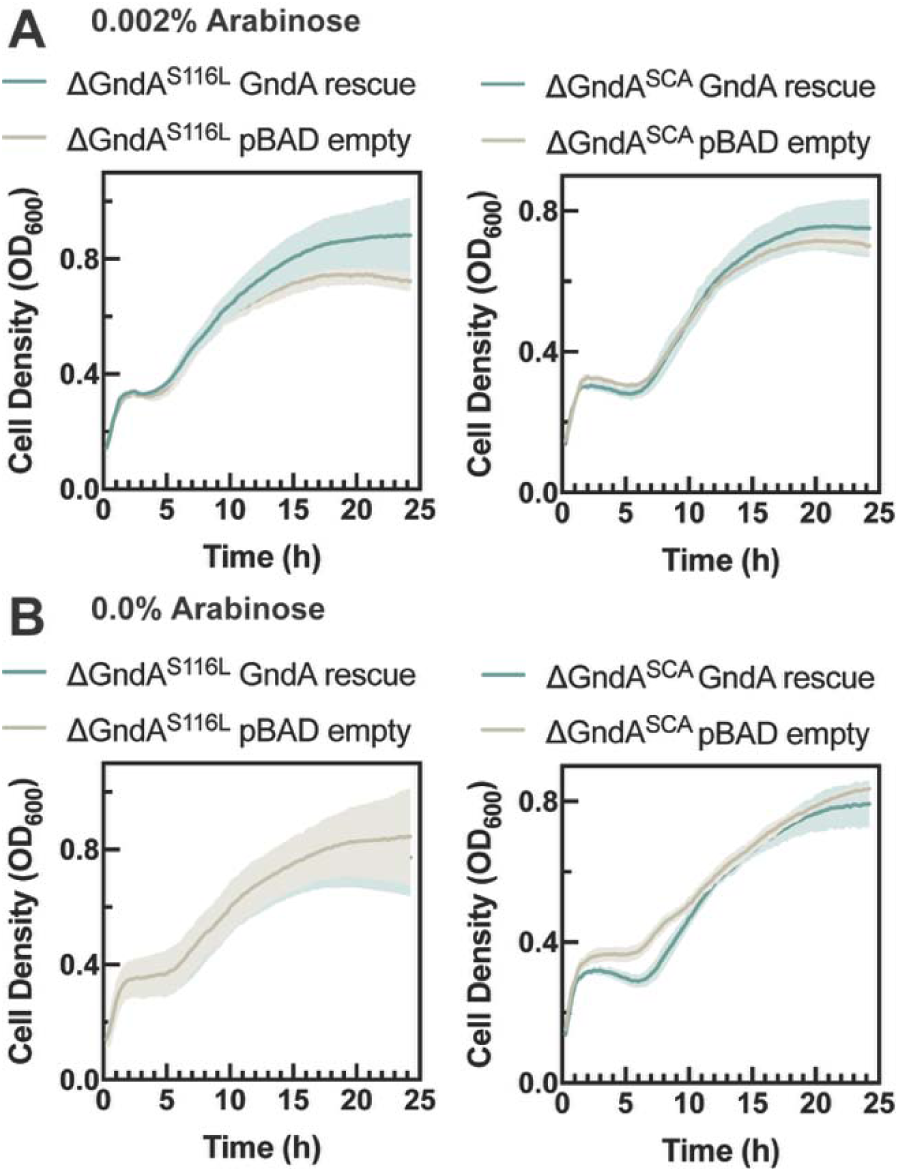
GndA reintroduction partially rescues growth of mutant strains at 45 °C. (A) ΔGndA^S116L^, left, and ΔGndA^SCA^, right, were transformed with a pBAD33 vector encoding GndA, or empty pBAD33 as a negative control. Growth curves were acquired at 0.002% arabinose and 45 °C. Error (shading above and below curves) calculated from 3 replicates. (B) The same strains were grown in the absence of arabinose at 45 °C as a control for the effect of arabinose-dependent GndA induction.

Many microproteins bind to and regulate the function of macromolecular complexes^39^. To identify candidate interaction partners of GndA, we co-immunoprecipitated the endogenously expressed, SPA-tagged GndA microprotein from lysates of GndA^SPA^ cells grown at both 37 °C and 45 °C. Enrichment was quantified relative to matched cell lysates exposed to anti-FLAG beads pre-incubated with 3xFLAG peptide antigen, which served as negative controls (Figure 3A-B). At both temperatures, the strongest enrichment observed was for subunits of the pyruvate dehydrogenase complex (cyan dots). We additionally observed enrichment of inner membrane complexes required for electron transport and ATP generation, including members of the *nuo* operon which encodes respiratory complex I (RCI) of the electron transport chain (red dots), as well as subunits of ATP synthase (black dots). To validate the protein enrichments detected with mass spectrometry, we performed co-immunoprecipitation and Western blotting. Subunits of candidate interacting complexes were appended with a myc epitope tag in an arabinose-inducible plasmid and transformed into the GndA^SPA^ strain. Anti-FLAG immunoprecipitation was performed from lysates of cells grown with arabinose induction, with anti-FLAG beads pre-incubated with 3xFLAG peptide antigen again serving as negative controls. SPA-GndA co-immunoprecipitated both the E1 subunit of the pyruvate dehydrogenase complex and the NuoCD subunit of RCI with very low background in the negative control (Figure 3C-D). Collectively, these findings suggest that endogenously expressed, SPA-tagged GndA associates with the pyruvate dehydrogenase complex and RCI in *E. coli*, either directly or indirectly. However, we cannot completely exclude the possibility that the association of GndA with inner membrane proteins may result from the insolubility of GndA, a phenomenon previously observed in the nonspecific enrichment of mitochondrial membrane proteins by mitochondrially localized human microproteins^19^. These results nonetheless likely reflect the subcellular environment in which GndA localizes and exerts its function.

**Figure 4.**
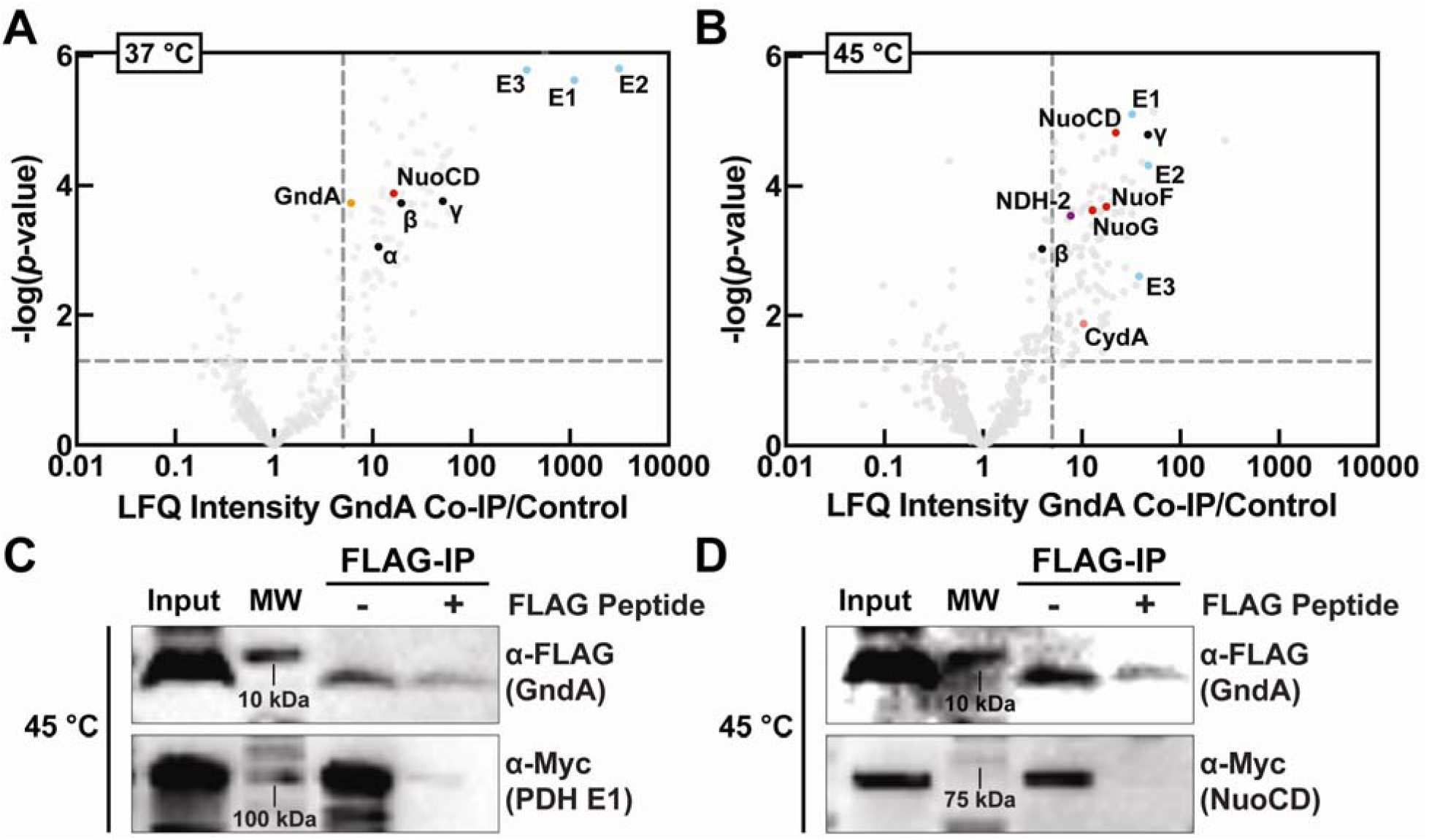
GndA co-immunoprecipitates with pyruvate dehydrogenase (PDH) and respiratory complex I (RCI). (A-B) Volcano plot of label free quantitative proteomic analysis of endogenous SPA-GndA co-immunoprecipitation from GndA^SPA^ cell lysates with reference to a preincubated 3xFLAG peptide control grown at (A) 37 °C and (B) 45 °C. *P*-values were calculated by performing a *t*-test assuming a two-tailed distribution and homoscedasticity (n = 4). Enriched subunits of PDH (blue), respiratory complexes (red) and ATP synthase (black) are highlighted. (C-D) Western blotting was carried out for co-immunoprecipitation of GndA with candidate interaction partners identified in (A-B). Myc epitope-tagged (C) pyruvate dehydrogenase subunit E1 or (D) respiratory complex 1 subunit NuoCD was expressed from an arabinose-inducible pBAD33 vector in GndA^SPA^ cells endogenously expressing SPA-tagged GndA. For each sample, a single cell lysate was divided into two tubes, from which anti-FLAG immunoprecipitation was carried out, with 3xFLAG peptide preincubated beads as a negative control. Anti-myc (PDH E1 or NuoCD, respectively) or anti-FLAG (GndA) Western blotting followed.

In order to maintain cellular homeostasis, gene regulatory networks are reshaped in response to stress and genetic perturbations. We therefore hypothesized that the transcriptional state of cells lacking GndA would provide additional insight into the role of GndA in the heat shock response. We analyzed differential gene expression via quantitative RNA sequencing (RNA-seq) of ΔGndA^SCA^ and ΔGndA^S116L^ strains relative to Δ*tolC* immediately after heat shock (Figure 4A-B). We observed that 119 genes were upregulated and 170 were specifically downregulated in both mutant strains after heat shock, demonstrating that GndA deletion and mutation result in similar changes to the *E. coli* gene expression program after heat shock. The top 5 upregulated genes in both datasets are metabolic and stress response proteins, including the toxic long direct repeat (ldr) peptide^40^ LdrC, while several of the most significantly downregulated genes in both datasets are involved in xylose utilization. After exclusion of the y-ome^32^, the top gene ontology (GO) term associated with genes upregulated in both mutant strains during heat shock is response to heat stress (Figure S5). These results clearly demonstrate that cells lacking GndA exhibit dysregulated expression of stress response and metabolic genes after heat shock, consistent with our observation of decreased fitness of these GndA mutants at high temperature (Figure 2A-B). We conclude that GndA contributes to maintenance of cellular homeostasis during *E. coli* growth at high temperature.

**Figure 5.**
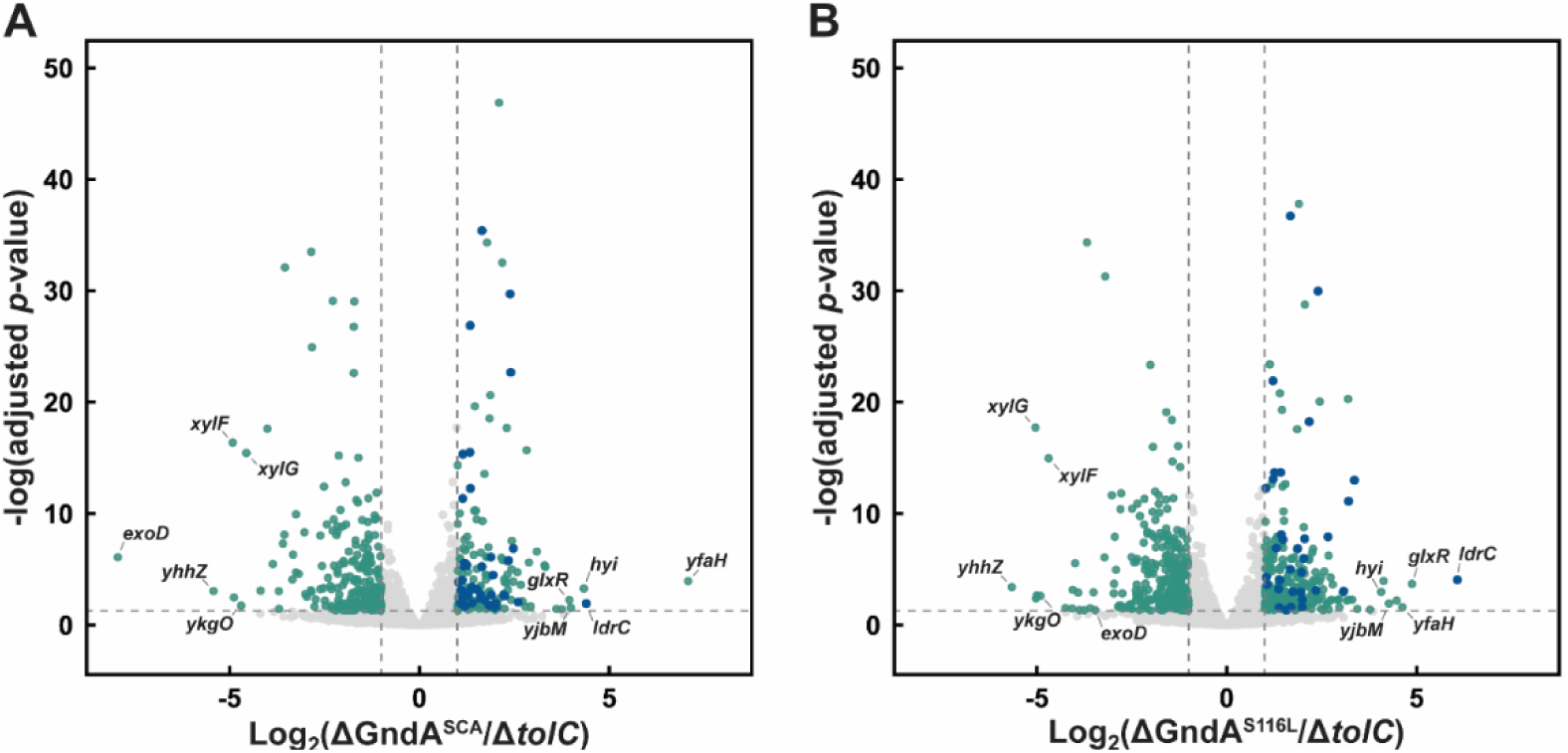
Loss of GndA alters the transcriptional response to elevated temperature. (*A*) RNA-seq analysis of gene expression changes in ΔGndA^SCA^ vs. Δ*tolC* and (*B*) ΔGndA^S116L^ vs. Δ*tolC* after growth at 45 °C. Adjusted *p*-values were calculated using DESeq2, which estimates differential expression using a negative binomial generalized linear model. *P*-values were derived from Wald tests and corrected for multiple testing using the Benjamini-Hochberg method (n = 3). Horizontal dotted line signifies *p*-value of 0.05, and vertical dotted lines signifies a fold change of 2. Blue dots signify up-regulated genes with functions in cellular stress responses. Green data points signify other significantly changing genes. The gene names of the 5 most significantly up- and down-regulated genes shared between both datasets are indicated.

## Discussion

In this work, we provide evidence for functionality of the 36-aa GndA microprotein in the bacterial heat shock response. The frameshifted *gndA* sORF internally overlaps the 6PGD CDS. While dozens of internal, overlapping ORFs like GndA have been identified in *E. coli* by ribosome profiling^6, 7^, very few have been characterized in any bacterial species to date (excepting *B. subtilis comS* and *T. thermophilus rpmH*). The study of internal ORFs poses unique challenges, as standard deletion strategies simultaneously perturb the overlapping protein CDS, leaving the functional significance of this class of sORFs largely unresolved. Our results thus provide direct evidence that an internal ORF can encode a functional microprotein.

GndA exhibits conservation in limited species, consistent with *de novo* origination and acquisition of functionality^5, 41^. While GndA is present in multiple gram-negative species, it is much less widely distributed than 6PGD, which is conserved from bacteria to eukaryotes. It is unclear in general how an internal ORF can arise *de novo* without creating deleterious mutations in the protein coding sequence that it overlaps. However, the *gnd* locus is unique in that it has previously been found to be polymorphic within *E. coli*, possibly due to its proximity to the O-antigen-encoding *rfb* region^42–47^. We therefore speculate that the polymorphic *gnd* locus is tolerant to mutations, which resulted in emergence of *gndA* in the +2 reading frame. A similar mechanism has previously been invoked for the emergence of overlapping viral genes within intrinsically disordered regions^48^. Notwithstanding its evolutionarily recent origination, the presence of GndA homologs in multiple species is consistent with our demonstration that it confers a fitness benefit during cellular stress.

Precise edits introduced in the GndA start codon or reading frame establish that loss of GndA leads to growth defects at 45 °C. While mutations in 6PGD also decreased cell fitness at high temperature, the growth defect observed in the GndA^SCA^ strain can be unambiguously attributed to loss of GndA. In addition, reintroduction of GndA on a plasmid into strains lacking it led to partial rescue of growth to varying degrees. Therefore, while both proteins encoded in the *gnd* locus support cell growth during heat shock, the GndA microprotein contributes a fitness benefit independent of 6PGD. While we focused in this work on establishing a role for GndA, we have not yet prepared or characterized isogenic strains lacking both GndA and 6PGD; further work will therefore be required to determine whether, like *comS* and *rpmH*, it regulates the same pathways or processes as the 6PGD protein that it overlaps – or not.

GndA may function in the heat shock response via its association with metabolic protein complexes. Endogenously expressed SPA-tagged GndA enriched the soluble pyruvate dehydrogenase complex as well as components of inner membrane-associated RCI from cell extracts at both 37 °C and 45 °C. It is not yet clear whether these interactions are direct or indirect. We therefore conclude that these protein associations reflect the biophysical properties and potential localization of GndA in cells. Future experiments will be required to test the hypothesis that GndA exerts its high-temperature fitness effect through its spatial or molecular associations.

Finally, we demonstrated that the transcriptional response to heat shock depends on GndA status, as two independent strains lacking GndA strongly upregulate stress response proteins relative to isogenic controls, demonstrating that *E. coli* experience more severe stress at elevated temperature in the absence of GndA. Collectively, these findings support a model in which GndA upregulation during heat shock suppresses the overexpression of toxic stress response proteins and promotes metabolic homeostasis and cell growth.

## Conclusion

This work establishes GndA as a functional microprotein that supports bacterial cell fitness during heat shock. This work furthermore provides genetic evidence for functionality of an internal, frameshifted sORF. More broadly, our findings expand the catalog of bacterial microproteins involved in stress responses and bioenergetics, challenge the assumption that small, poorly conserved ORFs lack biological relevance, and motivate the need to apply precision editing to uncharacterized internal ORF-encoded microproteins to determine if they are broadly functional.

## Data Availability

Proteomics data have been deposited to the ProteomeXchange Consortium via the PRIDE partner repository with the dataset identifier PXD065696.

RNA-seq data have been deposited to the GEO repository with accession number GSE330026.

## Acknowledgments

We thank all current and former members of the Slavoff and Breaker labs for helpful discussions. This work was funded in part by the National Institutes of Health (1R01GM155404-01), a Mark Foundation for Cancer Research Emerging Leader Award, a Paul G. Allen Frontiers Group Distinguished Investigator Award, and a Sloan Research Fellowship (FG-2022-18417) (to S.A.S.); and a NIGMS Chemistry-Biology Interface Training Program (1T32GM149444 to K.J., N.B. and C.D.). J.J.M. was supported in part by a Yale University fellowship associated with a NIGMS Chemistry-Biology Interface Training Program (5T32GM067543) and a Roberts Fellowship from the Yale University Department of Chemistry. M.G.M. is a Howard Hughes Medical Institute (HHMI) awardee of the Life Sciences Research Foundation and received supplementary support from HHMI funds awarded to the Breaker lab. C.D. was supported in part by a Gruber Science Fellowship from the Gruber Foundation. G. D. was supported in part by a National Science Foundation Graduate Research Fellowship (DGE-2632154). Any opinions, findings, and conclusions or recommendations expressed in this material are those of the authors(s) and do not necessarily reflect the views of the National Institutes of Health or National Science Foundation.

## Supporting Information

GndA conservation in Gram-negative bacteria; growth curves for wild-type and Δ*tolC* control strains, transformation and arabinose-induction controls at 37 °C; Gene Ontology analysis of the heat shock transcriptional response; oligodeoxynucleotide sequences and bacterial strain genotypes used in this study (Tables S1–S2, Figures S1–S5) (PDF)

## For Table of Contents Only

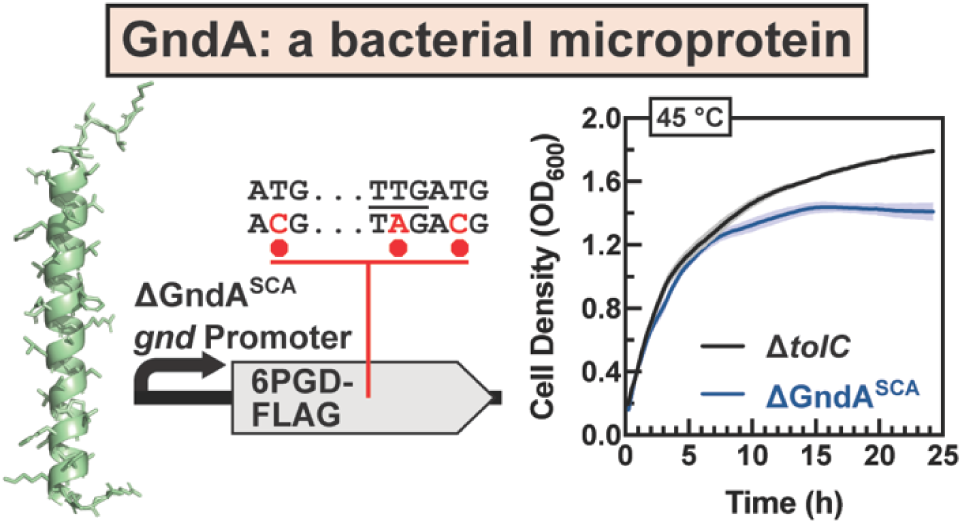

## Notes

https://www.ncbi.nlm.nih.gov/geo/query/acc.cgi?acc=GSE330026

### Competing Interest Statement

The authors have declared no competing interest.

### Summary of Updates

Figure 2 revised; new figure 3 added; figure 4 revised; supplementary information revised

