## Supplementary Information for "The GndA microprotein promotes cell growth during heat shock in *Escherichia coli*"

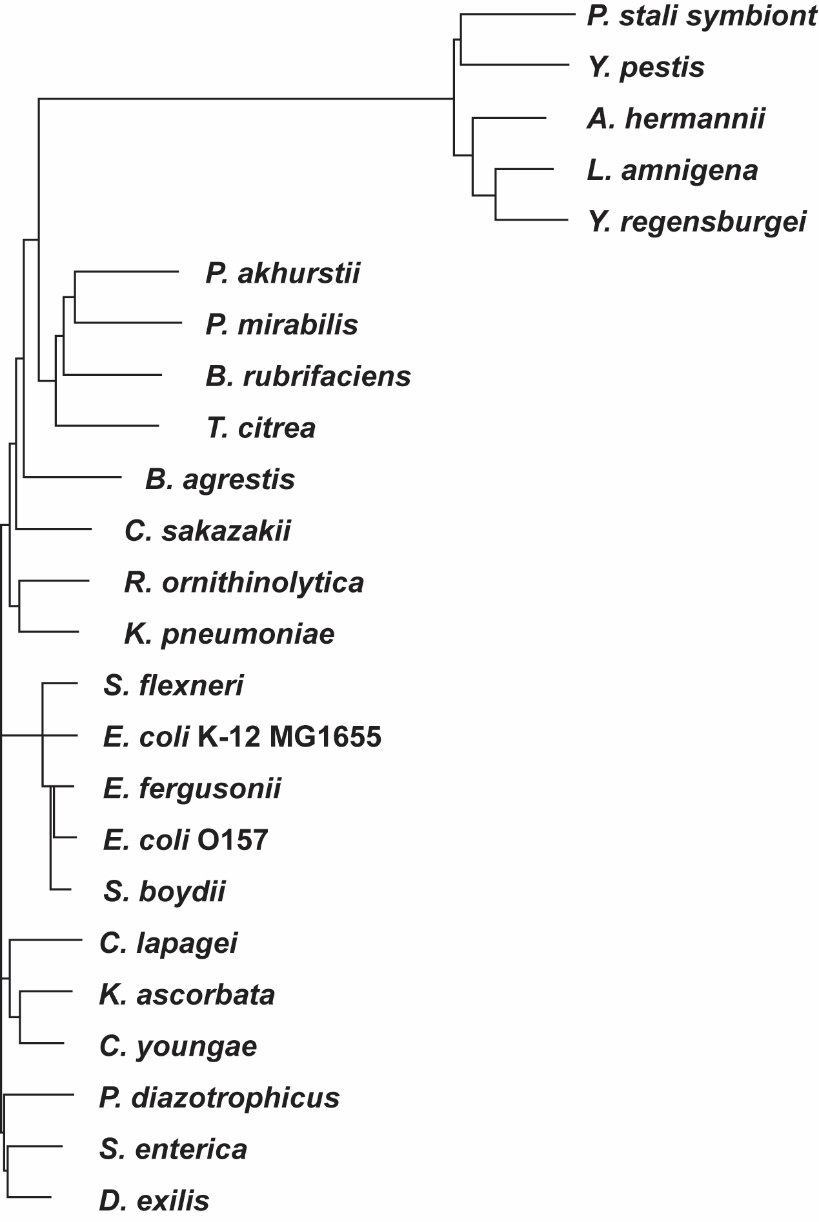

Figure S1. GndA conservation. 24 species of Gram-negative bacteria encoding putative syntenic GndA homologs were selected by tblastn (5) search of the *E. coli* K-12 MG1655 GndA sequence, and sequences exhibiting > 50% amino acid similarity were retained. The nucleic acid sequence of the putative homolog was inspected for the presence of a start and stop codon and Shine-Dalgarno consensus sequence. Alignment and phylogenetic tree for the nucleic acid sequences were constructed with Clustal Omega.

### Table S1. List of Custom Oligodeoxynucleotides.

| Name | nt | Sequence (5′→3′) |
| --- | --- | --- |
| C108 FLAG-GndA Fwd Primer MCS1pETDuet-1 | 60 | AAAAACATGCCATGGACTACAAAGACGATGACGACAAGGTGGTAACACCTTCTTCCAGGA |
| C108 FLAG-GndA Rev Primer MCS1pETDuet-1 | 33 | TTTTCCGGAATTCTCAGCGCCCCCTCTTCACCG |
| C108 NuoJ-cMyc Fwd Primer MCS2pETDuet-1 | 39 | AAAGGGAATTCCATATGGAGTTCGCTTTTTATATCTGTG |
| C108 NuoJ-cMyc Rev Primer MCS2pETDuet-1 | 66 | TTTTCCGCTCGAGTCAAAGATCTTCTTCGCTAATAAGTTTTTGTTCTGCGTGCTCCTCCGTTTTTC |
| *tolC* KO Oligo | 90 | A*A*ATGTGAATTTCAGCGACGTTTGACTGCCGTTTGAGCAGTCATGTGTTAAAGCTTCGGCCCCGTCTGAACGTAAGGCAACGTAAAGATA |
| Δ*gnd* Fwd Primer | 70 | TCATCAGTTTTTCACCCGTAATATAAAGCCGTAAGCATATAAGCATGGATTTGAGGCACATTAACGCCCT |
| Δ*gnd* Rev Primer | 68 | GCCCGGTGCAATATACGCCGGGCCTCAATTTTATTGTTGGTTAAATCAGAGTCTTCCGGGACCAGTGG |
| Δ6PGD Fwd Primer | 50 | TCATCAGTTTTTCACCCGTAATATAAAGCCGTAAGCATATAAGCATGGAT |
| Δ6PGD Rev Primer | 20 | GCCCGGTGCAATATACGCCG |
| ΔGndA Fwd Primer | 31 | TCATCAGTTTTTCACCCGTAATATAAAGCCG |
| ΔGndA Rev Primer | 20 | GCCCGGTGCAATATACGCCG |

### Table S2. Genotypes of *E. coli* K-12 MG1655 cell lines constructed in this study.

| Strain Name | Genotype |
| --- | --- |
| WT | Wild-type *E. coli* K-12 MG1655 |
| Δ*tolC* | Δ*tolC* |
| Δ*gnd* | Δ*gnd* (ΔGndA, Δ6PGD) |
| Δ6PGD | Δ6PGD, FLAG-GndA (epitope tag inserted after Thr9), Δ*tolC* |
| ΔGndA^S116L^ | ΔGndA, 6PGD-FLAG (S116L), Δ*tolC* |
| ΔGndA^A117V^ | ΔGndA, 6PGD-FLAG (A117V), Δ*tolC* |
| ΔGndA^SCA^ | ΔGndA, 6PGD-FLAG, Δ*tolC* |
| 6PGD^A148V^ | 6PGD-FLAG (A148V), Δ*tolC* |
| GndA^SPA^ | SPA-GndA, Δ6PGD, Δ*tolC* |

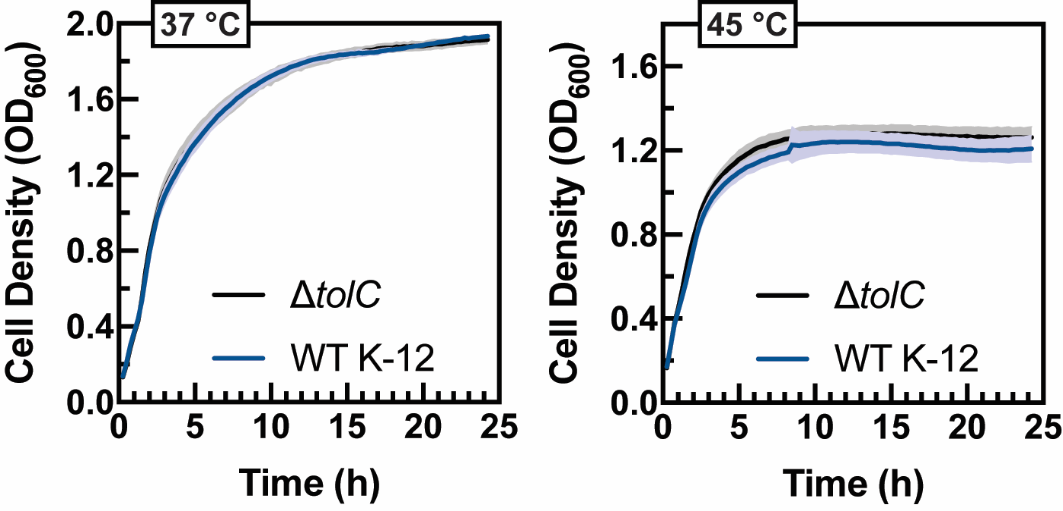

Figure S2. Comparison of wild-type *E. coli* K-12 and Δ*tolC* strain growth at 37 and 45 °C. Growth curves (OD_600_) for Δ*tolC* mutant (black) and WT K-12 (blue), grown at the indicated temperatures. Error (shading above and below curves) calculated from 3 replicates.

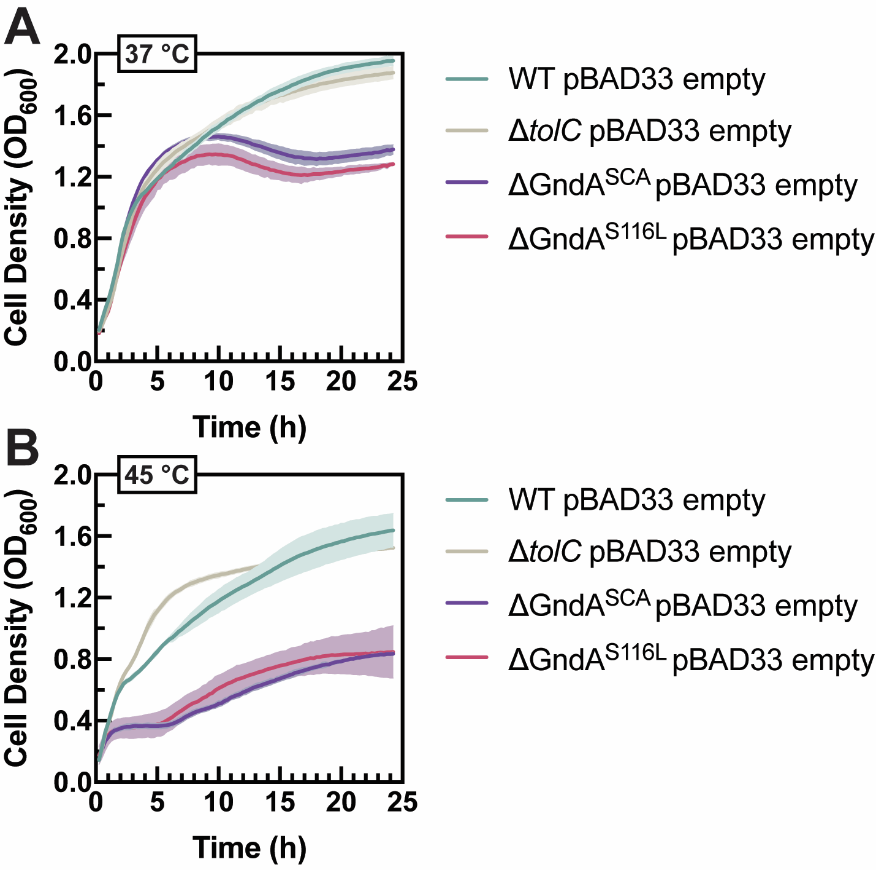

Figure S3. Growth of wild-type K-12, Δ*tolC*, ΔGndA^SCA^ and ΔGndA^S119L^ *E. coli* strains after plasmid transformation. The four *E. coli* K-12 MG1655 wild-type and mutant strains were freshly transformed with empty pBAD33 vector and grown at (A) 37 °C and (B) 45 °C in the absence of arabinose. Error (shading above and below curves) calculated from 3 replicates.

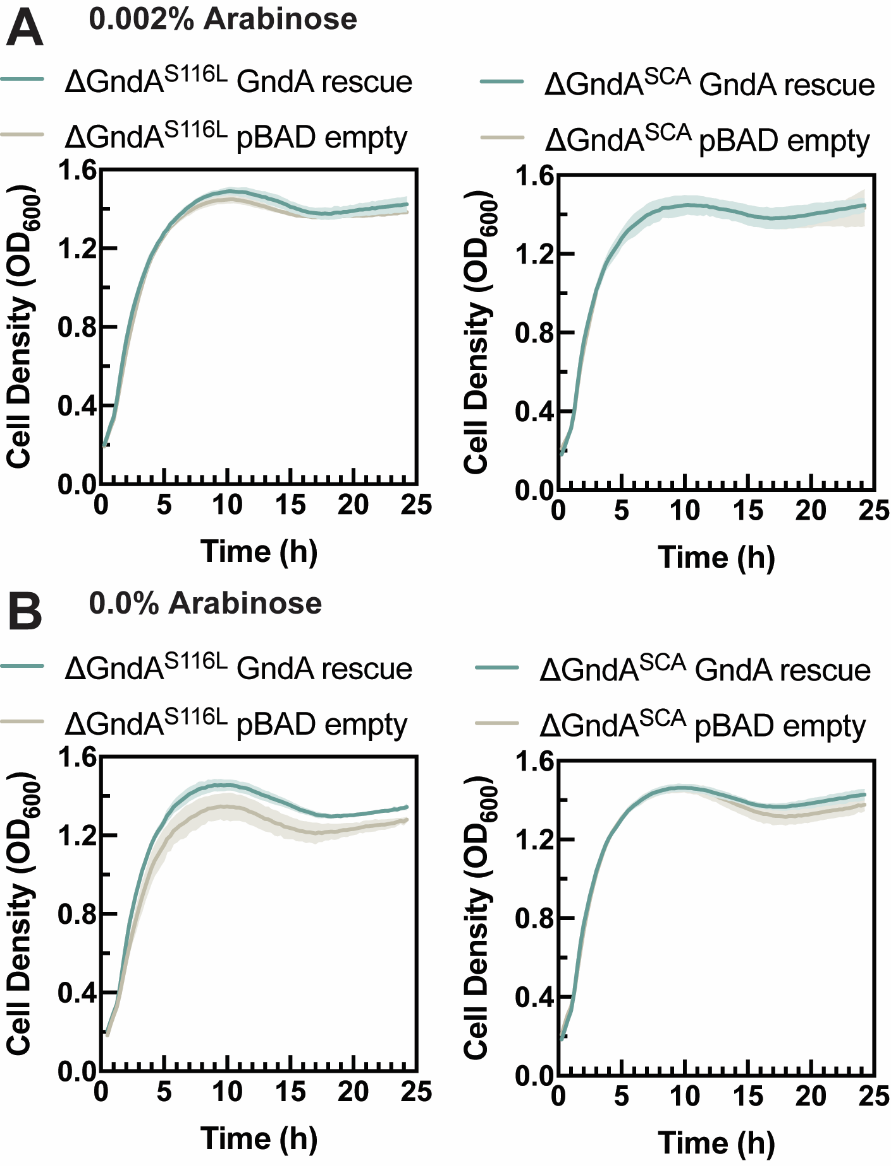

Figure S4. GndA rescue experiment at 37 °C. (A) ΔGndA^S116L^, left, and ΔGndA^SCA^, right, were transformed with a pBAD33 vector encoding GndA, or empty pBAD33 as a negative control. Growth curves were acquired at 0.002% arabinose and 37 °C. Error (shading above and below curves) calculated from 3 replicates. (B) The same strains were grown in the absence of arabinose at 37 °C as a control for the effect of arabinose-dependent GndA induction.

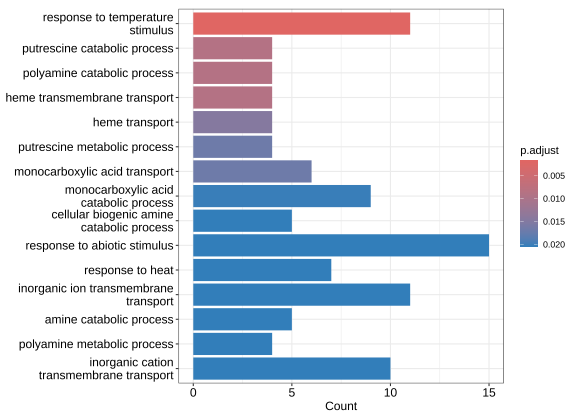

Figure S5. Gene Ontology (GO) analysis of the dysregulated transcriptional response to high temperature in two independent *E. coli* strains lacking GndA. RNA-seq analysis of differential gene expression in ΔGndA^SCA^ vs. Δ*tolC* and ΔGndA^S116L^ vs. Δ*tolC* after growth at 45 °C was quantified with DESeq2. Genes upregulated under these conditions in both mutant strains were subjected to GO analysis.
